# Transposon mobilization in the human fungal pathogen *Cryptococcus deneoformans* is mutagenic during infection and promotes drug resistance *in vitro*

**DOI:** 10.1101/2020.01.29.924845

**Authors:** Asiya Gusa, Jonathan D. Williams, Jang-Eun Cho, Anna Floyd-Averette, Sheng Sun, Eva Mei Shouse, Joseph Heitman, J. Andrew Alspaugh, Sue Jinks-Robertson

## Abstract

When transitioning from the environment, pathogenic microorganisms must adapt rapidly to survive in hostile host conditions. This is especially true for environmental fungi that cause opportunistic infections in immunocompromised patients since these microbes are not well adapted human pathogens. *Cryptococcus* species are yeast-like fungi that cause lethal infections, especially in HIV-infected patients. Using *Cryptococcus deneoformans* in a murine model of infection, we examined contributors to drug resistance and demonstrated that transposon mutagenesis drives the development of 5-fluoroorotic acid (5FOA) resistance. Inactivation of target genes *URA3* or *URA5* primarily reflected the insertion of two transposable elements (TEs): the T1 DNA transposon and the TCN12 retrotransposon. Consistent with *in vivo* results, increased rates of mutagenesis and resistance to 5FOA and the antifungal drugs rapamycin/FK506 and 5-fluorocytosine (5FC) were found when *Cryptococcus* was incubated at 37° compared to 30° *in vitro*, a condition that mimics the temperature shift that occurs during the environment-to-host transition. Inactivation of the RNAi pathway, which suppresses TE movement in many organisms, was not sufficient to elevate TE movement at 30° to the level observed at 37°. We propose that temperature-dependent TE mobilization in *Cryptococcus* is an important mechanism that enhances microbial adaptation and promotes pathogenesis and drug resistance in the human host.

**SIGNIFICANCE STATEMENT:** The incidence of infections due to fungal pathogens has dramatically increased in the past few decades with similar increases in human populations with weakened or suppressed immune systems. Understanding the mechanisms by which organisms rapidly adapt during human infection to enhance virulence and evolve drug resistance is important for developing effective treatments. We find that transposon mobilization in the human pathogen *Cryptococcus* causes genomic mutations in a murine model of infection and promotes resistance to antifungal drugs *in vitro*. Thermotolerance is a key virulence determinant for pathogenic fungi during the environment-to-host transition, and we demonstrate that a temperature increase is sufficient to trigger transposon mobilization *in vitro*. The link between temperature stress and transposon-associated mutations may significantly impact adaptation to the host during infection, including the acquisition of drug resistance.

## INTRODUCTION

*Cryptococcus* species (1) are environmental fungi that can cause lethal infections in highly immunocompromised patients. However, exposure to these fungal species is almost universal in many parts of the world, infecting humans by the inhalation of airborne spores from a variety of environmental sources such as soil and trees. Once inhaled, *Cryptococcus* can persist in a dormant, latent form, often only resulting in symptomatic infections in individuals with a suppressed or weakened immune system (2). *C. neoformans* (serotype A) is the predominant cause of HIV/AIDS-associated cryptococcal meningitis, which accounts for 15% of AIDS-related deaths and disproportionately impacts sub-Saharan Africa (3). *C. deneoformans* (serotype D) is more prevalent in Europe where it infects nearly 20% of HIV-positive patients (4) and is more commonly associated with skin lesions (5). *C. gattii* (serotypes B and C), which is endemic in tropical climates, is largely responsible for cryptococcal infections in immunocompetent hosts (2), including an ongoing outbreak in the more temperate region of the Pacific Northwest in North America (6).

*Cryptococcus* has several characteristics that contribute to its virulence and persistence in the human host. A dense polysaccharide capsule allows the yeast to evade host immune detection and escape phagocytosis by macrophages (7). Production of the cell wall-associated pigment melanin additionally protects against oxidative stress and temperature stress (8). Finally, the ability of *Cryptococcus* to withstand high temperatures, such as those found in mammalian hosts, is a defining feature of disease-causing fungi (9). *C. neoformans*, which is more virulent than *C. deneoformans*, is also more heat-resistant (10), consistent with thermotolerance as a key virulence factor.

During human infection, pathogenic fungi encounter environmental stresses, host immune defenses and treatment with antifungal drugs. In response, these fungi can undergo rapid genomic adaptations that enhance survival and enable drug resistance (11). Previously reported alterations arising during cryptococcal infection include base substitutions, insertions, deletions, target gene copy-number variation, and chromosomal duplications (12-14). These microevolutionary changes have been documented in AIDS patients with chronic and recurrent cryptococcal infections (12-16), as well as in murine models of infection (13, 16). Resulting phenotypes include resistance to azole drugs (12, 14) and differences in capsule structure, melanization and thermotolerance (12, 13, 17, 18), all of which are factors that increase persistence of *Cryptococcus* in the host. In addition, up to 20% of cryptococcal cells form giant polyploid cells (titan cells) during infection that are resistant to phagocytosis and have enhanced survival and dissemination (19, 20).

To further study the types of spontaneous genetic changes that arise in *Cryptococcus* during the stress of a host infection, a forward-mutation assay was used to identify 5FOA-resistant mutants in mice infected with *C. deneoformans* strain XL280α (21). We found that the majority of mutations resulting in drug-resistance were due to gene inactivation by transposable elements (TEs). Additional analyses *in vitro* demonstrated that TE-mediated mutagenesis was frequent during growth at a higher temperature that mimicked host infection (37°), but rarely occurred at the temperature normally used during laboratory growth (30°). The temperature dependence of transposition-associated drug resistance was confirmed in several clinical *C. deneoformans* strains and extended to clinically relevant antifungal drugs. We suggest that transposon mobilization activated by temperature stress is a previously uncharacterized adaptation mechanism that contributes to virulence and acquired drug resistance of *Cryptococcus* during infection.

## RESULTS

The spontaneous genetic alterations (i.e. base substitutions, insertions, deletions) that occur in *C. deneoformans* during infection were investigated by intravenously infecting eight mice with independently incubated inocula of strain XL280α. Fungi were recovered from the lungs, kidneys, and brain of each mouse between four and ten days post-infection, and mutants resistant to 5FOA were selected. Resistance to this drug occurs through inactivating mutations in either the *URA3* or *URA5* gene. We compared the frequency and types of spontaneous *URA3* and *URA5* mutations among the 5FOA-resistant mutants present in the inoculum cultures with those identified after passage in mice.

Mice were inoculated with ∼3×10^5^ cryptococcal cells and we recovered 10^7^ to 10^8^ cells per mouse, indicating a minimum of six to ten replication cycles within the host (*SI Appendix*, Table S1). For most mice, the fungal burden was highest in the kidneys and lowest in the brain (*SI Appendix*, Table S1). A total of 335 5FOA-resistant mutants were recovered from the eight mice, with a median mutant frequency of 6.9×10^−7^ per mouse (*SI Appendix*, Table S2). It should be noted that 5FOA-resistant mutant frequencies measured in mice may be an underestimate, as the uracil auxotrophy (Ura^-^ phenotype) associated with 5FOA-resistance in *C. neoformans* results in attenuated virulence in macrophage, *Galleria* and mouse models of infection (22). Interestingly, the median mutant frequency in cells recovered from the lungs (2.5×10^−6^) was ∼4-fold higher than that in the inoculum cultures or the kidneys; there were too few cells isolated from the brain to estimate a frequency (*SI Appendix*, Table S2). This difference could reflect a higher incidence of mutations arising in the lungs and/or more robust growth of Ura^-^ mutants in lung versus kidney tissue.

Given the frequency of 5FOA-resistance in the inoculum cultures and the number of cells used in each infection, it is unlikely that pre-existing 5FOA-resistant mutants were injected into most mice (median of 0.17 mutants per inoculum)(*SI Appendix*, Table S2). This leads to the prediction that most of the mutations identified in the post-passage cultures arose during the period of mouse infection. The *URA3* and *URA5* genes were thus PCR-amplified from genomic DNA of the 335 5FOA-resistant mutants to define the molecular nature of causative mutations. This analysis revealed the presence of 3 to 5 kb insertions within the *URA3* or *URA5* gene in 80% (269/335) of mutants recovered from mouse organs (Fig. 1*A*, *SI Appendix*, Table S3). Similar insertions were identified when *Cryptococcus* was grown in culture at 30° and 37° (Fig. 1*B*, described below). Sequencing of the larger-than-expected PCR amplicons identified insertions of either the DNA transposon Thing1 (T1; Genbank accession # AY145841) or the retrotransposon TCN12 (Genbank accession # JQ277268) (Fig. 1*C*). An additional 20 mutants had smaller insertions in *URA3/URA5* that corresponded to the 150-bp long terminal repeat (LTR) of TCN12 (Fig. 1*A*). Such solo LTRs presumably reflect deletional recombination between the LTR direct repeats of TCN12, as has been documented for the Ty1 retrotransposon of *Saccharomyces cerevisiae* (23). The remaining mutant isolates from which wildtype-sized *URA3* and *URA5* alleles were amplified likely contained base substitutions or small insertions/deletions.

**Figure 1.**
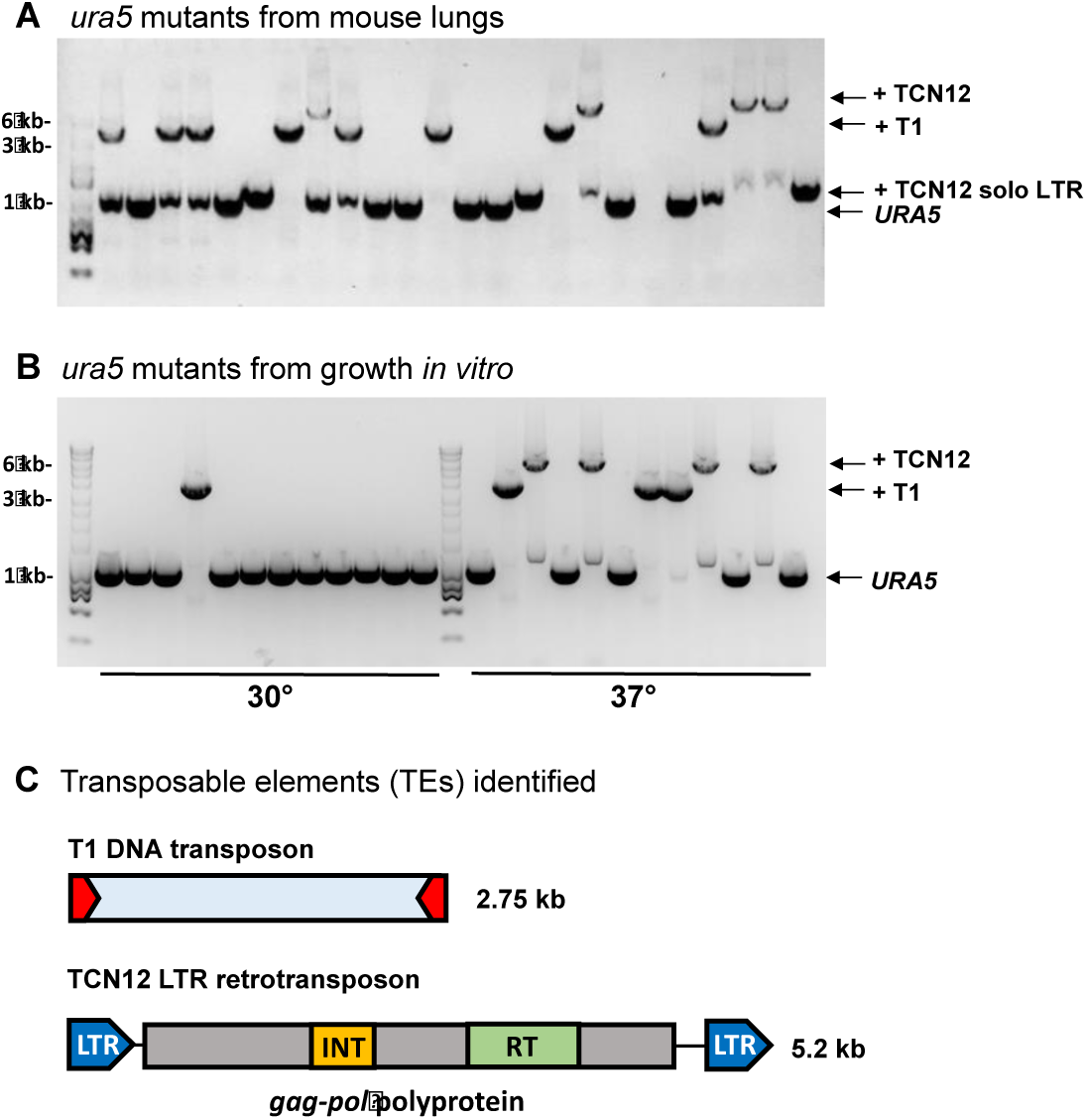
PCR-amplification of *URA5* in representative 5FOA-resistant mutants showing T1, TCN12 (full-length), and TCN12 solo LTR (long terminal repeat) insertions. *(A)* Amplification from genomic DNA of mutants recovered from the lungs of Mouse 3. *(B)* Amplification from genomic DNA of mutants isolated after growth at 30° or 37° *in vitro. (C)* Illustration of the T1 DNA transposon with 11-bp terminal inverted repeats and the TCN12 retrotransposon with the *gag-pol-*polyprotein and 150-bp LTR direct repeats. The integrase (INT) and reverse transcriptase (RT) open reading frames are indicated. Amplification of a wildtype-sized *URA5* band in addition to the larger band diagnostic of TE insertion presumably reflects loss of the TE in a subpopulation of cells during non-selective growth prior to isolation of genomic DNA.

Figure 2 summarizes the identity, location, and orientation of each unique T1 and TCN12 insertion identified in 5FOA-resistant mutants in inoculum cultures and after passage in mice. Strains recovered from mouse 3 had the largest number of unique insertions; these as well as those identified in the corresponding inoculum culture are highlighted in gray. As expected, insertion of these TEs duplicated the target-site sequence, creating short direct repeats flanking the element. Target-site duplication size is characteristic of each element and corresponds to the distance between the integrase-generated nicks that allow TE insertion (24). Insertion of T1 was associated with 3-bp duplications while that of TCN12 generated 5-bp target-site duplications (underlined in Fig. 2).

**Figure 2.**
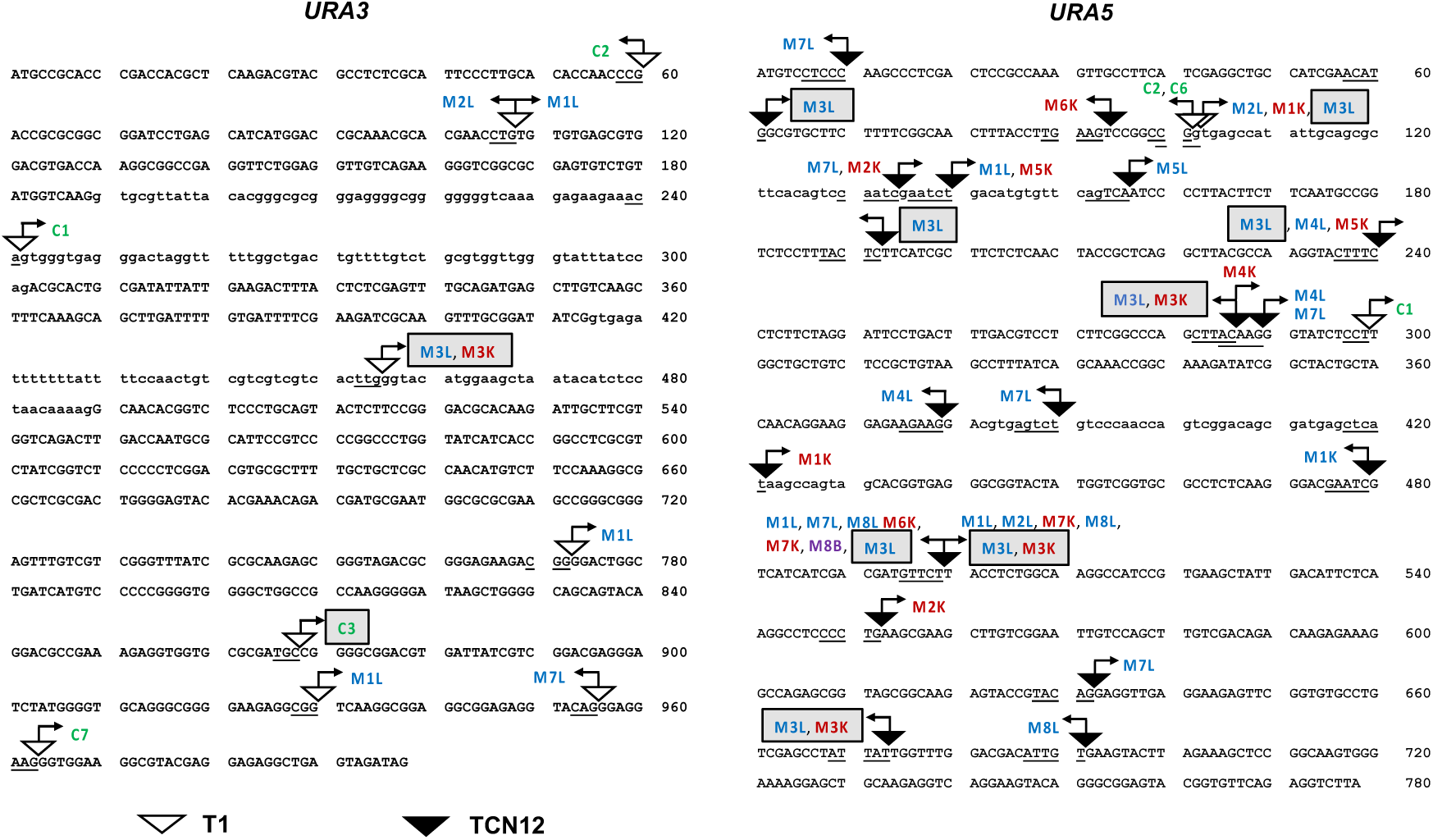
TE insertions (T1 and TCN12) in *URA3* and *URA5* of 5FOA-resistant mutants recovered from mice (M1-M8; color coded by organ) and inoculum cultures (C1-C7; green font). C7 was used to infect both M7 and M8. Highlighted in gray are TE insertions from mouse 3 and culture 3. The organ from which each mutant was recovered is indicated: lungs (L) blue, kidneys (K) red, brain (B) purple. Arrows represent the forward or reverse integration of each TE. Uppercase letters are exon sequences, lowercase letters are intron sequences. Underlined sequences indicate those sequences that are duplicated following TE integration. For some of the TCN12 insertions, only a solo LTR was detected.

Some TE insertions were recovered multiple times from mice or the inoculum cultures, suggesting they were clonally derived (*SI Appendix*, Table S3). Importantly, however, *none* of the insertions recovered from mice were identical to those detected in the inocula (Fig. 2). In total, insertional inactivation by TEs (full-length or solo LTR) accounted for 86% of mutations recovered from mice, with 43 unique insertion events among the 335 mutants detected (Fig. 2, *SI Appendix*, Table S3). By contrast, a smaller proportion of mutants screened from the inoculum cultures (56/168) had TE insertions in *URA3* or *URA5*, and there were only seven unique TE events among 168 mutants analyzed (Fig. 2, *SI Appendix*, Table S3). Comparing the numbers of unique TE insertions recovered from mice to the number found in the 30° inoculum cultures, significantly more insertions arose during infection (p=0.002 by Fisher exact test)(*SI Appendix*, Table S3).

One striking feature of the data was the locus specificity of TCN12 (Fig. 2). Although the numbers are small, T1 insertions appeared to be equally distributed between *URA3* and *URA5* (p=0.45 by Chi-square goodness of fit) and were found in inocula as well as after mouse passage. By contrast, TCN12 insertions were detected only after passage through mice. Additionally, among 34 independent TCN12 insertions identified (those found in different organs from the same mouse were not considered independent), *all* were in *URA5*. These data suggest that conditions in the mouse host uniquely stimulated TCN12 movement, with a very strong bias for insertion into *URA5*. Finally, there was evidence of target-site specificity for both T1 and TCN12. Of 16 independent T1 insertions, two occurred after base pair position 108 in *URA3*, two occurred after position 101 in *URA5*, and three occurred after position 102 in *URA5*. In the case of TCN12, of the 34 independent insertions, three insertions occurred after position 240, two after position 286, two after position 289, and ten after position 499 in *URA5* (Fig. 2).

### Temperature-dependent TE mobilization *in vitro*

Because growth temperature was a key difference between 5FOA-resistant *Cryptococcus* mutants recovered from inocula (30°) and those recovered from mice (∼37°), we investigated whether the mobilization of TEs can be triggered by temperature stress. In this analysis, XL280α was incubated in non-selective medium (YPD) for three days at 30° or 37° prior to selective plating on drug-containing media. Growth at 37° increased the rates of resistance to 5FOA, rapamycin/FK506 and 5-fluorocytosine (5FC) between 7.8- and 28-fold depending on the drug (Fig. 3*A*).

**Figure 3.**
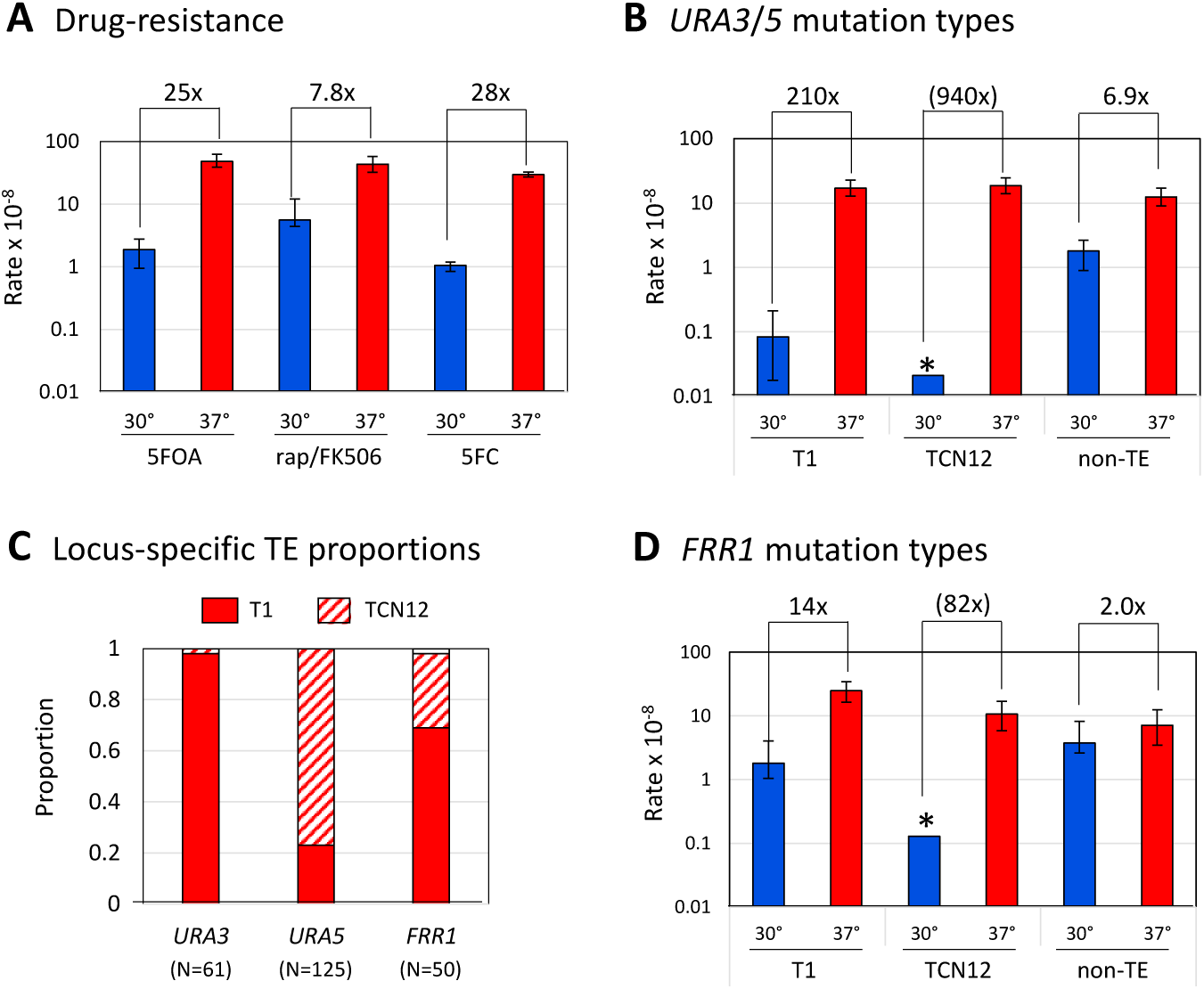
Temperature-dependent mutagenesis is driven by TE insertions. (*A*) Drug-resistance rates for 5FOA, rapamycin/FK506 (rap/FK506) and 5-fluorocytosine (5FC) when growth before drug selection was at either 30° (blue) or 37° (red) *in vitro*. (*B*) Insertion rates of individual TEs into *URA3* or *URA5 (URA3/5).* (*C*) Distribution of T1 and TCN12 insertions in *URA3, URA5* and *FRR1* in drug-resistant mutants isolated from 37° cultures; the distribution at each locus is significantly different (p<0.001, Chi-square contingency test). (D) Insertion rates of T1 and TCN12 into the *FRR1* gene. Error bars are 95% confidence intervals; rates are statistically different if error bars do not overlap. Asterisks indicate no insertions were detected; the rate shown was calculated assuming one event.

In the case of 5FOA-resistant mutants that arose during *in vitro* growth, only those derived from different cultures were analyzed in order to ensure independence. Similar to *in vivo* results following mouse infection, PCR analysis of the *URA3* and *URA5* genes revealed T1 or TCN12 insertions in ∼75% (186/251) of mutants that arose during growth at 37° (Fig. 1*B*, *SI Appendix*, Fig. S1, Table S4). By contrast, only ∼5% (4/90) of the mutants arising at 30° amplified *URA3* and *URA5* alleles of larger-than-expected size (p<0.0001). We estimate that the rates of T1 and TCN12 insertions were elevated more than 200-fold during growth at 37°compared to 30° (Fig. 3*B*). 5FOA-resistant mutants that produced wildtype-sized *URA3* and *URA5* amplification products were assumed to reflect base substitution or small insertions/deletions. Although not further investigated here, the rates of non-TE events were also significantly elevated at the higher temperature, but not to the level of TE insertions (Fig. 3*B*).

Among 251 independent 5FOA-resistant mutants isolated following *in vitro* growth at 37°, there were 61 TE insertions into *URA3* and 125 into *URA5*. In *URA3*, 60 of the 61 TE insertions were T1, with only one TCN12 insertion detected. At *URA5*, however, there were 96 insertions of TCN12 and 29 insertions of T1 (Fig. 3*C*, *SI Appendix*, Fig. S1, Table S4). In terms of locus preference, there was a 2-fold bias for T1 insertion into *URA3* (p=0.001 by Chi-square goodness of fit) and an estimated 100-fold bias for TCN12 insertion into *URA5* (p<0.0001). In addition, there was a strong hotspot for TCN12 insertion into *URA5*, with 46% (44/96) of insertions occurring after position 499. This is the same hotspot of TCN12 insertion found among the 5FOA-resistant mutants recovered from mice (Fig. 2). In the case of T1, there were no more than three insertions at a given position within *URA3* or *URA5*.

In addition to examining the nature of mutations conferring 5FOA resistance, we examined whether TE elements contributed to rapamycin/FK506 resistance, which occurs through inactivation of the *FRR1* gene encoding FKBP12 (25). Similar to 5FOA-resistance, TE insertions were the major cause of increased rapamycin/FK506 resistance at 37° (Fig. 3*D*). As in the 5FOA assay, TCN12 transposition was only detected at 37°; T1 insertion occurred at both temperatures but was elevated more than 10-fold at 37° (Fig. 3*D*, *SI Appendix*, Table S4). Additionally, a single insertion of the DNA transposon Thing2 (T2; Genbank accession #AY145844) into *FRR1* was detected at each temperature (*SI Appendix*, Table S4).

The combination of antifungal drugs amphotericin B and 5FC is an effective treatment for cryptococcosis (26). 5FC-resistance in fungi can result from inactivation of *FCY1* (a cytosine deaminase), *FCY2* (a cytosine permease) or *FUR1* (a uracil phosphoribosyltransferase)(27, 28). Mutation in *UXS1*, a capsule biosynthesis enzyme, was additionally identified as causative of 5FC-resistance in *C. deuterogattii* (28). In a screening of independent 5FC-resistant XL280 mutants obtained *in vitro*, we identified T1 and TCN12 insertions in the *UXS1* gene (*SI Appendix*, Fig. S2). T1 insertions were found in mutants arising at 30°, and both T1 and TCN12 insertions were found in mutants arising at 37°, consistent with the temperature-dependent TE profile in 5FOA- and rapamycin/FK506-resistant mutants. Because a complete set of causative mutations mediating 5FC-resistance in *Cryptococcus* is not yet known (28), we were unable to assess the rate of 5FC drug-resistance in XL280 due to TE insertions.

### Temperature-dependent inactivation of RNAi is not the cause of TE mobilization

RNA interference (RNAi) has been shown to restrain movement of some TEs in *Cryptococcus* species (29-31), consistent with its role in many other systems (32). To examine whether the observed temperature-dependent TE mobilization was simply the result of RNAi inactivation at 37°, we deleted the *RDP1* gene, which encodes the RNA-dependent RNA polymerase (RdRP) that synthesizes double-stranded RNAs in the RNAi pathway (33). If temperature-dependent inactivation of RNAi is the primary cause of TE mobilization, then the rate of TE movement at 30° in the *rdp1*Δ strain should be similar to that in the wildtype strain at 37°. At 30°, the 5FOA-resistance rate increased ∼2.5 fold in the *rdp1*Δ background, and the proportion of T1 insertions increased from 4/90 to 23/54 (*SI Appendix*, Table S4). Although this corresponded to a 26-fold increase in T1 insertions, the rate was still 8.5-fold lower than that in the *RDP1* strain when grown at 37° (Fig. 4). In the *rdp1*Δ background no movement of TCN12 was detected at 30°, and the rate of TCN12 insertions was not further elevated at 37°. We note that non-LTR retrotransposon T3 (also known as CNIRT4; Genbank accession # KC440173.1), whose movement was not detected in the 341 5FOA-resistant mutants isolated in the *RDP1* background, was inserted into *URA3/URA5* in 6/121 mutants isolated in the *rdp1*Δ strain (p=0.0003 by Fisher exact test)(*SI Appendix*, Table S4). Although these results confirm that RNAi limits the movement of some TEs in *Cryptococcus*, inactivation of RNAi potentially makes only a small contribution to the temperature-dependent mobilization of T1 or TCN12 observed here.

**Figure 4.**
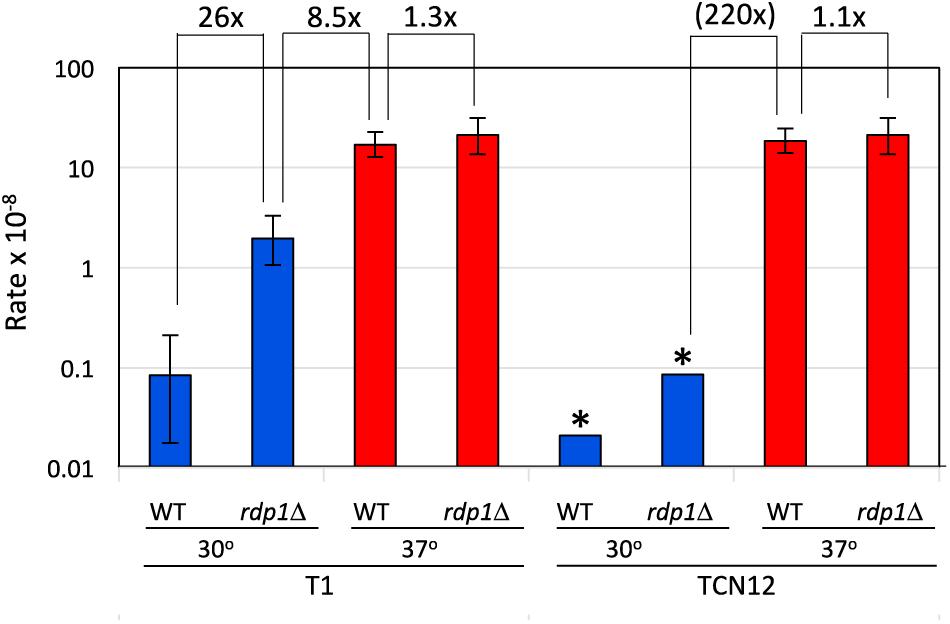
RNAi is not responsible for temperature-dependent control of TE movement. Insertion rates of T1 and TCN12 into *URA3/5* in wild-type (WT) and RNAi-deficient (*rdp1*Δ) strains grown at 30° (blue) or 37° (red). Error bars are 95% confidence intervals; rates are statistically different if error bars do not overlap. Asterisks indicate no insertions were detected; the rate shown was calculated assuming one event.

### TE mobilization occurs in both clinical and environmental isolates of *C. deneoformans*

XL280 is a laboratory strain derived from crosses between an environmental (NIH433) and a clinical (NIH12) isolate (34), and it is virulent in mice (35). To determine if temperature-dependent transposon mobilization also occurs in the ancestral strains, 5FOA-resistance rates were measured following growth at 30° and 37°. We also analyzed two additional clinical isolates, 528 and AD7-71 (36), to determine if this phenomenon extends to other *C. deneoformans* strains. In strains NIH12, NIH433, and 528, the 5FOA-resistance rate was elevated 4- to 6-fold at 37°; there was no significant effect of temperature on mutation rate in strain AD7-71 (Fig. 5*A* and *SI Appendix*, Table S4). Further molecular analysis revealed that only clinical strains NIH12 and 528 displayed increased TE insertions into *URA3/URA5*; no TE insertions were detected in NIH433 at either temperature (*SI Appendix* Table S4). T1 insertions were modestly elevated at 37° in both NIH12 and 528 (*SI Appendix*, Table S4). By contrast, TCN12 insertions were only detected in NIH12 and were elevated 38-fold at the higher temperature (Fig. 5*B*). As observed in XL280, TCN12 insertions were strongly biased to the *URA5* locus (18/19 insertions). Insertion of T3, previously only detected in RNAi-deficient XL280 (29) (*SI Appendix*, Table S4), was common in both strains and also was elevated at 37° (Fig. 5*B*). These data indicate that the temperature-dependent TE mobilization in XL280 was inherited from its clinical progenitor and that other clinical isolates share this general phenotype. The temperature-dependent mobilization of specific TEs, however, varies from strain to strain.

**Figure 5.**
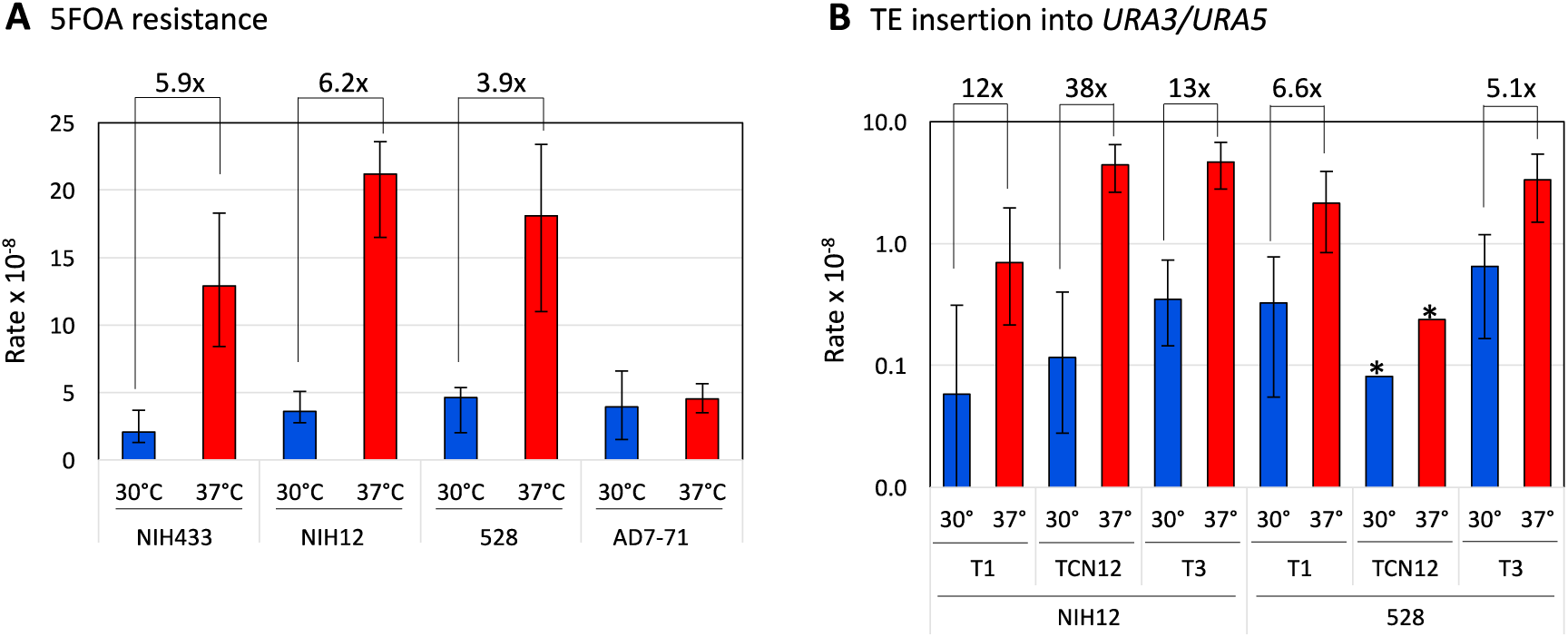
5FOA resistance rates and TE insertion rates in clinical and environmental isolates. (*A*) 5FOA resistance rates for clinical strains NIH12, 528, AD7-71, and environmental strain NIH433 grown at 30° (blue) or 37° (red). The ratios of the rates at the two temperatures are indicated by the brackets. (*B*) Insertion rates of TEs into *URA3/5* in clinical isolates NIH12 and 528. Error bars are 95% confidence intervals; rates are statistically different if error bars do not overlap.

## DISCUSSION

In this study, we report the active mobilization of transposable elements as a significant driver of mutagenesis in *C. deneoformans* during mouse infection and growth at elevated temperatures *in vitro* (Figs. 1-3). Transposon-mediated inactivation of target genes was shown to confer drug resistance to 5FOA and clinically relevant antifungal drugs rapamycin/FK506 and 5FC (Fig. 3; SI Appendix, Fig. S2). We found that TE mobilization was increased dramatically at the host-relevant temperature of 37° compared to 30° *in vitro* (Fig. 3), strongly linking the activation of TE movement to heat stress. Importantly, this temperature-dependent TE movement also was observed among several clinical strains surveyed (Fig. 5), indicating that the phenomenon is not limited to the XL280 lineage. Mutagenesis in response to heat stress during infection likely contributes to phenotypic variation that promotes adaptation, survival and virulence as well as the development of antifungal-drug resistance. Further, the current warming of environmental temperatures may foster a more rapid adaptation to higher, host-relevant temperatures and enhanced pathogenesis. If similar, temperature-triggered TE mutagenesis extends to other non-pathogenic fungal species, it may facilitate the breaching of the temperature barrier that currently prevents them from causing disease in human populations and other mammals (37).

Transposable elements make up a significant portion of eukaryotic genomes, comprising ∼14% in fungi, 45 to 50% in human cells and up to 85% in plants (32, 38, 39). Since McClintock’s discovery of TEs in the 1940s (40), the impact of mobile genetic elements on gene expression, gene regulation, and genome rearrangement has been the focus of ongoing studies (38). TEs play both passive and active roles in shaping genomes and driving evolutionary change. In terms of rearrangements, multiple copies of a given element can serve as substrates for non-allelic homologous recombination that creates inversions, deletions, translocations and complex rearrangements (11). Such TE-mediated rearrangements are thought to occur in all organisms including fungi, plants, and humans (38). In *Cryptococcus*, where 5% of the genome is comprised of TEs (41), transposons have been implicated in facilitating gene rearrangements and contributing to the divergence and speciation between *C. neoformans* and *C. deneoformans* (42).

In *Cryptococcus*, we demonstrated that drug resistance to 5FOA, rapamycin/FK506 and 5FC increased significantly at 37° *in vitro* (Fig. 3). Drug resistance at elevated temperature resulted primarily from insertional inactivation of target genes *URA3* and *URA5* (5FOA resistance) and *FRR1* (rapamycin/FK506 resistance) (Fig. 3). Additionally, we identified TE insertions in one of the gene targets mediating 5FC resistance (*UXS1*)(*SI Appendix*, Fig. S2). Since *uxs1* mutants are deficient in capsule synthesis and avirulent in a mouse model of infection (43), it is unclear whether these mutants would propagate during infection with or without 5FC treatment. A screening of known target genes from 5FC clinical isolates or from resistant isolates following mouse infection would be useful to determine if transposon mutagenesis is causative of drug resistance during infection. The forward mutation strategy used in this study was only able to detect TE insertion in a very small segment (∼5 kb) of the *Cryptococcus* genome. In addition, the clear target-site specificity of TCN12 suggests that there may have been elements whose movement we failed to detect. While the current study focused on the inactivation of specific genetic targets that conferred drug resistance, a more global analysis of temperature-dependent TE mobilization will likely reveal aspects of target-site specificity in terms of local chromatin structure, for example, and may reveal the mobilization of additional elements.

Gene disruption leading to loss of protein function, as used here, is a major outcome of random transposition, but insertions can also modulate nearby gene expression. For example, insertion of TEs into promoter regions can upregulate or downregulate gene expression by altering access of trans-acting regulatory proteins, or can introduce cis-elements in promoter and enhancer regions that lead to changes in gene expression (38). In the case of retrotransposons such as TCN12, the LTR-associated promoter can also constitutively activate nearby gene expression. Fluconazole resistance, for example, is a major problem in the treatment of cryptococcal infections and is mediated by increased expression of target genes (*ERG11/AFR1*)(44). Although characterized drug-resistant mutants typically have chromosomal duplications that increase target gene copy number and gene expression (44), transposon insertions such as those reported may also contribute to acquired drug resistance by affecting local gene expression.

The cryptococcal genome is remarkably plastic under the stressful conditions of host infection and this can lead to drug resistance (11, 45). Because the movement of transposons can cause chromosomal breakage and disruption of gene function, transcriptional and posttranscriptional mechanisms have evolved to suppress TE movement and limit mutagenic effects (32). These mechanisms include RNAi silencing to target dsRNA for degradation and epigenetic alterations (for example, DNA methylation) of genomic sequences to limit TE transcription (32). *Cryptococcus* has a functional RNAi silencing system that previously was found to suppress the movement of DNA transposons T2 and T3 as well as the retrotransposon Cnl1 (29). In this study, we detected increased mobility of the T1 and T3 transposons, but not of TCN12, in an RNAi-defective *rdp1*Δ derivative of XL280 at 30° (Fig. 4, *SI Appendix*, Table S4), confirming that RNAi silencing regulates the movement of these transposons. Their mobility at 30° in the absence of *RDP1*, however, was still much lower than at 37° in the presence of *RDP1*, and deletion of *RDP1* did not further elevate movement at 37°. An as-yet undescribed mechanism was thus largely responsible for the temperature-dependent mobilization described here.

Movement of the TCN12 retrotransposon in *Cryptococcus* was dramatically increased both at elevated temperature in culture and in the mouse model of infection (Figs. 1-3). Indeed, TCN12 movement was only detected at elevated temperatures (Fig. 3). Heat-induced transposition has been studied most extensively in plants and *Drosophila* (46, 47). In *Arabidopsis thaliana*, movement of the *ONSEN* retrotransposon is coupled with the heat stress response (48). This LTR-retrotransposon contains heat-responsive element (HRE) sequences in its promoter, which are recognized and activated by heat shock factor A 2 (HSFA2) as part of the plant’s general response to temperature stress (46, 48). A similar mechanism of transcriptional control in response to heat stress is found in other, copia-like LTR retrotransposons in *Brassicaceae* species (46). Whether TCN12 is controlled at the transcriptional level or through a post-transcriptional mechanism is not yet known.

Darwin hypothesized that evolution does not occur at a constant rate, but that dramatic adaptive changes occur in relatively short bursts (49). TE mobilization in response to environmental stresses is one such mechanism of rapid and potentially adaptive mutagenesis and has been well documented in plants and *Drosophila* (50-52). Our results further strengthen this model by demonstrating that a fungal pathogen undergoes rapid TE-mediated genomic alterations that are triggered by conditions in a mammalian host. This additional, TE-mediated genetic diversity in *Cryptococcus* may contribute to evasion of the host immune system and the development of novel virulence-promoting phenotypes, as well as to the emergence of drug resistance. Further studies are needed to explore the global impact of transposon movements throughout the cryptococcal genome and to obtain a better understanding of the identity, prevalence and mechanism of heat-responsive transposable elements in *Cryptococcus* and other fungal genomes.

## MATERIALS AND METHODS

### Strains and growth media

XL280α (21) is a *C. deneoformans* strain derived by the subsequent crossing of progeny spores of a diploid obtained by mating environmental strain NIH433 and clinical strain NIH12 (34, 53). Clinical strains 528 and AD7-71 were from European patients (36). Cells were grown non-selectively in YPD medium (1% yeast extract, 2% Bacto-peptone, 2% dextrose; 2% agar for plates). Mutants resistant to 5FOA were selected on synthetic complete medium (SC; 0.17% yeast nitrogen base, 0.5% ammonium sulfate, 0.13% Hartwell’s complete amino acid mix and 2% agar) supplemented with 1.3 μg/mL uracil and 1 μg/mL 5FOA (US Biological). Rapamycin/FK506-resistant mutants were selected on YPD supplemented with 1 μg/mL of rapamycin (LC Laboratories) and 1 μg/mL FK506 (Astellas). For 5FC selection, cells were plated on pre-mixed synthetic complete medium (Becton Dickinson) supplemented with 100 μg/mL 5FC.

Two independent *RDP1* mutants were generated in the XL280α background by separate biolistic transformation of a PCR-generated *rdp1Δ::NAT* cassette amplified from the genomic DNA of an XL280 strain previously used to characterize RNAi pathways (29). The forward and reverse primers used for PCR were 5’GCATGATTTGCATCCATGAATCCCGCCATGTAGACTTCC and 5’GTCTCACTACTCATACTTGTCAAAGATGGTACCTTCGAGAAGCTGAGG, respectively, and LA Taq (Takara Bio) was used for amplification. Following selection of nourseothricin-resistant transformants on drug-supplemented YPD, deletion of *RDP1* was confirmed by PCR.

### Mouse infection and mutant recovery

Independent colonies of XL280α grown on non-selective YPD medium for two days at 30° were inoculated into 100 mL YPD and incubated at 30° overnight with shaking. 50 mL of each culture was pelleted and washed three times with phosphate-buffered saline (PBS). Cell concentration was determined using a hemocytometer and cells were diluted to a final concentration of 10^7^ cells/mL for mouse infections. An aliquot of each cell suspension was plated on selective 5FOA and non-selective YPD media at 30° to select for 5FOA-resistant mutants in the inoculum and to verify the infection dose, respectively.

Ten female A/J mice (6-8 weeks old) were used in this study (Jackson Labs). Seven mice (1-7) were each infected intravenously by retroorbital injection with 50 μL (target dose of 5 ×10^5^ cells) of independent XL280α cell suspensions. An additional mouse (mouse 8) was infected with the same cell suspension as mouse 7 but with a 5-fold lower dose of cells (1×10^5^). Two mice were injected with 50 μL PBS as controls. Mice were monitored daily for signs of illness such as weight loss, fatigue and loss of equilibrium. Mice died or were sacrificed by CO_2_ inhalation between days 4 and 10 post-infection. The lungs, kidneys and brain of each infected mouse were harvested immediately after CO_2_ inhalation or within 12 hours of death and organs were homogenized separately in 1 mL PBS. Each homogenate was plated on 5FOA plates (200 μL on each of four plates) to select for mutants resistant to 5FOA. The remaining homogenate was diluted and plated non-selectively on YPD to determine the cryptococcal burden. Kanamycin (50 μg/mL) and chloramphenicol (30 μg/mL) were added to plates to prevent bacterial growth and plates were incubated at 30°.

### Mutation rate determination *in vitro*

Each culture was inoculated with a colony grown for two days at 30° on YPD medium. Cells were grown non-selectively in YPD liquid medium on a roller drum at either 30° or 37°, as appropriate, and at least 24 independent cultures were used for each rate determination. After three days of growth, cells were pelleted, washed and resuspended in water. The total numbers of cells and drug-resistant mutants in each culture were determined by plating appropriate dilutions on YPD and drug-containing plates, respectively. With the exception of rapamycin/FK506 selection, which required incubation at 37°, all other drug selections and non-selective growth were at 30°. Colonies were counted four days after selective plating and mutation rates were calculated using the method of the median (54). The 95% confidence interval (CI) for each rate was determined as previously described (55).

### Mutation Spectra

Following genomic DNA isolation, candidate genes (*URA3* and *URA5* for 5FOA resistance; *FRR1* for rapamycin/FK506 resistance; and *UXS1* for 5FC resistance) were screened by PCR using MyTaq polymerase (Bioline). Forward (F) and reverse (R) primers were F-5’GATCCCGCTGATTAGTGGAGTCG and R-5’GTATCACCATGCTTATTGCGTATTC for *URA3*; F-5’CGCGTACGGCGATGTCCGCC and R-5’CACGCCTGCCTGTACTTAAGTTC for *URA5*; F-5’CTATCCGGACGTGCAATACTTTCC and R-5’CTTTAAGACCATCTACTAATTTGCACAATAAG for *FRR1*; and F-5’AGTGCCCAGGCACGGAAAGAAGAAGCTC and R-5’CGCATCATCATTCCCGTCGCAAAATCAACTCG for *UXS1*. If amplification using MyTaq failed, PCR was repeated using the long-range polymerase LA Taq and primers with a higher melting temperature. Primers used were F-5’GATCCCGCTGATTAGTGGAGTCGAACCACACCAG and R-5’GTATCACCATGCTTATTGCGTATTCATATCAACTATGCGTG for *URA3*; F-5’CGCGTACGGCGATGTCCGCCTCCAC and R-5’CACGCCTGCCTGTACTTAAGTTCCTTTGATACAAGC for *URA5*; and F-5’GTCCGTCACCCCTATCCGGACGTGCAATACTTTCC and R-5’CTTTAAGACCATCTACTAATTTGCACAATAAGTTTAGTTGATCTTGAGG for *FRR1*. Products containing an insertion were sequenced from each end (Sanger sequencing, Eton Bioscience) to identify the junctions of the inserted element. The insertion rate was calculated by multiplying the proportion of mutants with a given TE by the overall drug-resistance rate. 95% CI for proportions were determined using vassarstats.net. To obtain a 95% CI for a given mutation type, a method employing the square root of the sum of the squares (using the component 95% CI for rates and proportions) was used (56).

## Acknowledgments

This research was supported by National Institutes of Health grants R35GM118077 and R21AI133644 to S.J.R, R01AI074677 to J.A.A., NIH/NIAID R01 AI50113-15 and R37 MERIT Award AI39115-21 to J.H. and the NIH Tri-Institutional Molecular Mycology and Pathogenesis Training Program postdoctoral fellowship (5T32AI052080 to J.D.W and 2T32AI052080 to A.G.). We thank Dr. Connie Nichols, Dr. Aaron Smith, and Calla Telzrow for their assistance in experiments.

## References

1. F. Hagen et al., Recognition of seven species in the *Cryptococcus gattii/Cryptococcus neoformans* species complex. Fungal Genet. Biol. 78, 16–48 (2015).

2. X. Lin, J. Heitman, The biology of the *Cryptococcus neoformans* species complex. Annu. Rev. Microbiol. 60, 69–105 (2006).

3. R. Rajasingham et al., Global burden of disease of HIV-associated cryptococcal meningitis: an updated analysis. Lancet Infect. Dis. 17, 873–881 (2017).

4. F. Dromer, S. Mathoulin-Pélissier, O. Launay, O. Lortholary, French Cryptococcosis study, determinants of disease presentation and outcome during cryptococcosis: the CryptoA/D study. PLoS Med. 4, e21 (2007).

5. F. Dromer et al., Individual and environmental factors associated with infection due to *Cryptococcus neoformans* serotype D. Clin. Infect. Dis. 23, 91–96 (1996).

6. C. A. D’Souza et al., Genome variation in *Cryptococcus gattii*, an emerging pathogen of immunocompetent hosts. mBio 2, e00342–00310 (2011).

7. A. Casadevall et al., The capsule of *Cryptococcus neoformans*. Virulence 10, 822–831 (2019).

8. D. Lee et al., Unraveling melanin biosynthesis and signaling networks in *Cryptococcus neoformans*. mBio 10, e02267–02219 (2019).

9. V. A. Robert, A. Casadevall, Vertebrate endothermy restricts most fungi as potential pathogens. J. Infect. Dis. 200, 1623–1626 (2009).

10. L. R. Martinez, J. Garcia-Rivera, A. Casadevall, *Cryptococcus neoformans* var. *neoformans* (serotype D) strains are more susceptible to heat than *C. neoformans* var. *grubii* (serotype A) strains. J. Clin. Microbiol. 39, 3365–3367 (2001).

11. A. Gusa, S. Jinks-Robertson, Mitotic recombination and adaptive genomic changes in human pathogenic fungi. Genes 10, 901 (2019).

12. Y. Chen et al., Microevolution of serial clinical isolates of *Cryptococcus neoformans* var. *grubii* and *C. gattii*. mBio 8, e00166–00117 (2017).

13. G. Hu et al., Variation in chromosome copy number influences the virulence of Cryptococcus neoformans and occurs in isolates from AIDS patients. BMC Genomics 12, 526 (2011).

14. N. R. H. Stone et al., Dynamic ploidy changes drive fluconazole resistance in human cryptococcal meningitis. J. Clin. Invest. 129, 999–1014 (2019).

15. D. Sullivan, K. Haynes, G. Moran, D. Shanley, D. Coleman, Persistence, replacement, and microevolution of *Cryptococcus neoformans* strains in recurrent meningitis in AIDS patients. J. Clin. Microbiol. 34, 1739–1744 (1996).

16. B. C. Fries, F. Chen, B. P. Currie, A. Casadevall, Karyotype instability in *Cryptococcus neoformans* infection. J. Clin. Microbiol. 34, 1531–1534 (1996).

17. D. Garcia-Hermoso, F. Dromer, G. Janbon, *Cryptococcus neoformans* capsule structure evolution in vitro and during murine infection. Infect. Immun. 72, 3359–3365 (2004).

18. B. C. Fries, A. Casadevall, Serial isolates of *Cryptococcus neoformans* from patients with AIDS differ in virulence for mice. J. Infect. Dis. 178, 1761–1766 (1998).

19. O. Zaragoza, K. Nielsen, Titan cells in *Cryptococcus neoformans*: cells with a giant impact. Curr. Opin. Microbiol. 16, 409–413 (2013).

20. L. H. Okagaki et al., Cryptococcal cell morphology affects host cell interactions and pathogenicity. PLoS Pathog. 6, e1000953–e1000953 (2010).

21. B. Zhai et al., Congenic strains of the filamentous form of *Cryptococcus neoformans* for studies of fungal morphogenesis and virulence. Infect. Immun. 81, 2626–2637 (2013).

22. F. A. de Gontijo et al., The role of the *de novo* pyrimidine biosynthetic pathway in *Cryptococcus neoformans* high temperature growth and virulence. Fungal Genet. Biol. 70, 12–23 (2014).

23. D. J. Garfinkel, K. M. Nyswaner, K. M. Stefanisko, C. Chang, S. P. Moore, Ty1 copy number dynamics in *Saccharomyces*. Genetics 169, 1845–1857 (2005).

24. H. H. Kazazian, Mobile elements: drivers of genome evolution. Science 303, 1626–1632 (2004).

25. M. C. Cruz et al., Rapamycin antifungal action is mediated via conserved complexes with FKBP12 and Tor kinase homologs in *Cryptococcus neoformans*. Mol. Cell. Biol. 19, 4101–4112 (1999).

26. M. S. Saag et al., Practice guidelines for the management of cryptococcal disease. Clin. Infect. Dis. 30, 710–718 (2000).

27. W. W. Hope, L. Tabernero, D. W. Denning, M. J. Anderson, Molecular mechanisms of primary resistance to flucytosine in *Candida albicans*. Antimicrob. Agents Chemother. 48, 4377–4386 (2004).

28. R. B. Billmyre, S. Applen Clancey, L. X. Li, T. L. Doering, J. Heitman, 5-fluorocytosine resistance is associated with hypermutation and alterations in capsule biosynthesis in *Cryptococcus*. Nat. Commun. 11, 127 (2020).

29. G. Janbon et al., Characterizing the role of RNA silencing components in *Cryptococcus neoformans*. Fungal Genet. Biol. 47, 1070–1080 (2010).

30. X. Wang et al., Sex-induced silencing defends the genome of Cryptococcus neoformans via RNAi. Genes Dev 24, 2566–2582 (2010).

31. X. Wang, S. Darwiche, J. Heitman, Sex-induced silencing operates during opposite-sex and unisexual reproduction in *Cryptococcus neoformans*. Genetics 193, 1163–1174 (2013).

32. R. K. Slotkin, R. Martienssen, Transposable elements and the epigenetic regulation of the genome. Nat. Rev. Genet. 8, 272–285 (2007).

33. E. Bernstein, A. A. Caudy, S. M. Hammond, G. J. Hannon, Role for a bidentate ribonuclease in the initiation step of RNA interference. Nature 409, 363 (2001).

34. X. Lin, C. M. Hull, J. Heitman, Sexual reproduction between partners of the same mating type in *Cryptococcus neoformans*. Nature 434, 1017–1021 (2005).

35. M. Feretzaki, S. E. Hardison, F. L. Wormley, Jr., J. Heitman, *Cryptococcus neoformans* hyperfilamentous strain is hypervirulent in a murine model of cryptococcal meningoencephalitis. PLoS One 9, e104432–e104432 (2014).

36. W. Li et al., Genetic diversity and genomic plasticity of *Cryptococcus neoformans* AD hybrid strains. G3: Genes, Genomes, Genetics 2, 83–97 (2012).

37. M. A. Garcia-Solache, A. Casadevall, Global warming will bring new fungal diseases for mammals. mBio 1, e00061–00010 (2010).

38. B. Chénais, A. Caruso, S. Hiard, N. Casse, The impact of transposable elements on eukaryotic genomes: From genome size increase to genetic adaptation to stressful environments. Gene 509, 7–15 (2012).

39. M.-J. Daboussi, P. Capy, Transposable elements in filamentous fungi. Ann. Rev. Microbiol. 57, 275–299 (2003).

40. B. McClintock, The origin and behavior of mutable loci in maize. Proc. Natl. Acad. Sci. U.S.A. 36, 344–355 (1950).

41. B. J. Loftus et al., The genome of the basidiomycetous yeast and human pathogen *Cryptococcus neoformans*. Science 307, 1321–1324 (2005).

42. S. Sun, J. Xu, Chromosomal rearrangements between serotype A and D strains in *Cryptococcus neoformans*. PLoS One 4, e5524 (2009).

43. F. Moyrand, B. Klaproth, U. Himmelreich, F. Dromer, G. Janbon, Isolation and characterization of capsule structure mutant strains of *Cryptococcus neoformans*. Molecular Microbiology 45, 837–849 (2002).

44. E. Sionov, H. Lee, Y. C. Chang, K. J. Kwon-Chung, *Cryptococcus neoformans* overcomes stress of azole drugs by formation of disomy in specific multiple chromosomes. PLoS Pathog. 6 (2010).

45. A. Forche, Large-scale chromosomal changes and associated fitness consequences in pathogenic fungi. Curr. Fungal Infect. Rep. 8, 163–170 (2014).

46. B. Pietzenuk et al., Recurrent evolution of heat-responsiveness in *Brassicaceae* COPIA elements. Genome Biol. 17, 209 (2016).

47. M. P. García Guerreiro, What makes transposable elements move in the *Drosophila* genome? Heredity (Edinb) 108, 461–468 (2012).

48. V. V. Cavrak et al., How a retrotransposon exploits the plant’s heat stress response for its activation. PLoS Genetics 10 (2014).

49. C. Darwin, On the Origin of Species, 1859 (Routledge, 2004).

50. B. McClintock, The significance of responses of the genome to challenge. Science 226, 792 (1984).

51. L. Piacentini et al., Transposons, environmental changes, and heritable induced phenotypic variability. Chromosoma 123, 345–354 (2014).

52. V. Specchia et al., Hsp90 prevents phenotypic variation by suppressing the mutagenic activity of transposons. Nature 463, 662–665 (2010).

53. M. Ni et al., Unisexual and heterosexual meiotic reproduction generate aneuploidy and phenotypic diversity *de novo* in the yeast *Cryptococcus neoformans*. PLoS Biol. 11, e1001653 (2013).

54. D. E. Lea, C. A. Coulson, The distribution of the numbers of mutants in bacterial populations. J. Genet. 49, 264–285 (1949).

55. R. M. Spell, S. Jinks-Robertson, Determination of mitotic recombination rates by fluctuation analysis in *Saccharomyces cerevisiae* in Genetic Recombination. (Springer, 2004), pp. 3–12.

56. A. Moore et al., Genetic Control of Genomic Alterations Induced in Yeast by Interstitial Telomeric Sequences. Genetics 209, 425–438 (2018).

