## Supplementary Appendix for "Transposon mobilization in the human fungal pathogen *Cryptococcus deneoformans* is mutagenic during infection and promotes drug resistance *in vitro*"

* Sue Jinks-Robertson.

**This PDF file includes:**

Figure S1 to S2

Tables S1 to S4

**SI Appendix**

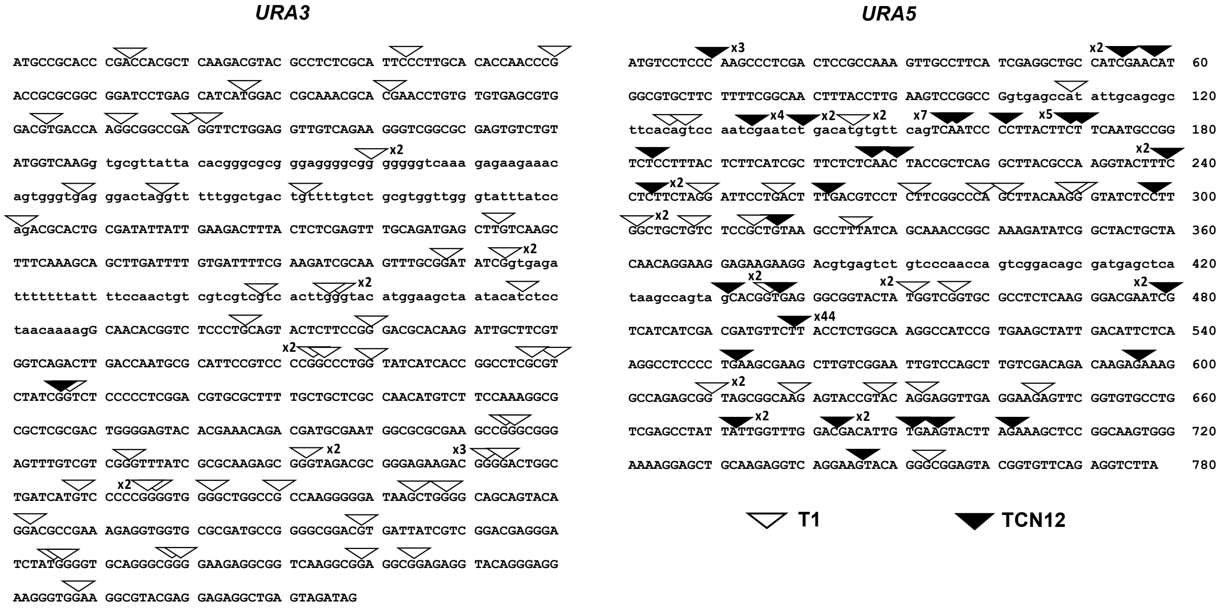


**Figure S1**. The numbers and locations of independent T1 and TCN12 insertions (white and black triangles, respectively) into *URA3* or *URA5* from cultures grown at 37°. Coding regions are represented by uppercase letters while intronic sequences are lowercase.

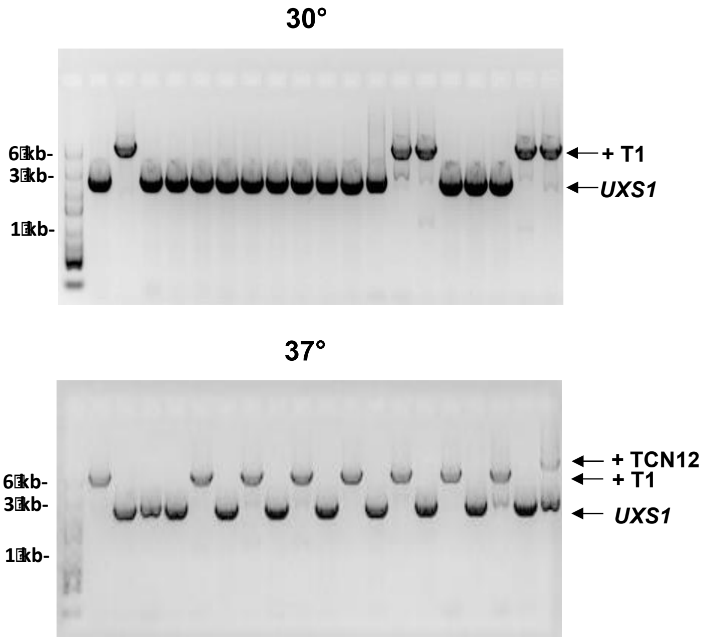


**Figure S2**. PCR-amplification of *UXS1* in representative 5-fluorocytosine (5FC) resistant mutants showing T1 and TCN12 insertions. Mutants were obtained after growth at 30° (top panel) or 37° (bottom panel) *in vitro*.

**Table S1.** Infection dose and number of *Cryptococcus* recovered from mouse organs.

| | MOUSE 1 | MOUSE 2 | MOUSE 3 | MOUSE 4 | MOUSE 5 | MOUSE 6 | MOUSE 7 | MOUSE 8 |
|---|---|---|---|---|---|---|---|---|
| Inoculum culture | 1 | 2 | 3 | 4 | 5 | 6 | 7 | 7 |
| Infection dose (cells/mouse) | $3.9x10^5$ | $5.6x10^5$ | $3.0x10^5$ | $2.9x10^5$ | $3.8x10^5$ | $3.1x10^5$ | $3.0x10^5$ | $6.1x10^4$ |
| Day organs harvested | DAY 4 | DAY 4 | DAY 5* | DAY 5* | DAY 6 | DAY 7 | DAY 10 | DAY 10 |
| Cells recovered from lungs | $1.5x10^7$ | $2.7x10^7$ | $2.6x10^7$ | $4.8x10^7$ | $2.6x10^7$ | $1.3x10^7$ | $2.2x10^7$ | $1.3x10^7$ |
| Cells recovered from kidneys | $2.5x10^7$ | $3.7x10^7$ | $7.2x10^7$ | $1.6x10^7$ | $6.9x10^7$ | $1.3x10^7$ | $5.2x10^7$ | $1.1x10^7$ |
| Cells recovered from brain | $2.0x10^6$ | $2.0x10^6$ | $8.5x10^5$ | $4.0x10^6$ | $2.3x10^6$ | $1.8x10^6$ | $6.6x10^6$ | $5.5x10^7$ |
| Total cells recovered from organs | $4.2x10^7$ | $6.6x10^7$ | $9.9x10^7$ | $6.8x10^7$ | $9.7x10^7$ | $2.8x10^7$ | $8.0x10^7$ | $7.9x10^7$ |
| Minimum # of generations | 6 | 6 | 8 | 8 | 6 | 6 | 7 | 10 |

*Mice died at 5 days post-infection, organs harvested within 12 hrs of death.

**Table S2.** Frequency of XL280 5FOA-resistant mutants in inoculum cultures and recovered from mice.

| INOCULUM | Culture 1 | Culture 2 | Culture 3 | Culture 4 | Culture 5 | Culture 6 | Culture 7 | |
|---|---|---|---|---|---|---|---|---|
| Number of mutants | 1741 | 277 | 135 | 7384 | 79 | 101 | 26 | |
| Mutant frequency | $2.9 \times 10^{-6}$ | $8.1 \times 10^{-7}$ | $5.6 \times 10^{-7}$ | $3.2 \times 10^{-5}$ | $1.9 \times 10^{-7}$ | $1.9 \times 10^{-7}$ | $2.7 \times 10^{-7}$ | |
| Estimated mutants in inoculum | 1.1 | 0.45 | 0.17 | 9.3 | 0.072 | 0.059 | 0.081 | |
| **MICE** | **MOUSE 1** | **MOUSE 2** | **MOUSE 3** | **MOUSE 4** | **MOUSE 5** | **MOUSE 6** | **MOUSE 7** | **MOUSE 8** |
| Inoculum culture | 1 | 2 | 3 | 4 | 5 | 6 | 7 | 7 |
| Day organs harvested | DAY 4 | DAY 4 | DAY 5 | DAY 5 | DAY 6 | DAY 7 | DAY 10 | DAY 10 |
| **LUNGS** | | | | | | | | |
| Number of mutants | 37 | 22 | 78 | 28 | 1 | 0 | 15 | 22 |
| Mutant frequency | $3.1 \times 10^{-6}$ | $1 \times 10^{-6}$ | $4.3 \times 10^{-6}$ | $2.5 \times 10^{-6}$ | $1.9 \times 10^{-7}$ | $<1.1 \times 10^{-7}$ | $2.5 \times 10^{-6}$ | $2.4 \times 10^{-6}$ |
| **KIDNEYS** | | | | | | | | |
| Number of mutants | 22 | 15 | 55 | 6 | 14 | 4 | 12 | 0 |
| Mutant frequency | $1.1 \times 10^{-6}$ | $5.0 \times 10^{-7}$ | $1.1 \times 10^{-6}$ | $1.4 \times 10^{-7}$ | $8.2 \times 10^{-7}$ | $4.4 \times 10^{-7}$ | $6.0 \times 10^{-7}$ | $<1.1 \times 10^{-7}$ |
| **BRAIN** | | | | | | | | |
| Number of mutants | 0 | 0 | 0 | 0 | 0 | 0 | 1 | 3 |
| Mutant frequency | $<6.3 \times 10^{-7}$ | $<7.1 \times 10^{-7}$ | $<1.7 \times 10^{-6}$ | $<1.0 \times 10^{-6}$ | $<2.4 \times 10^{-6}$ | $<7.7 \times 10^{-7}$ | $2.2 \times 10^{-7}$ | $6.8 \times 10^{-8}$ |
| **ALL ORGANS** | **MOUSE 1** | **MOUSE 2** | **MOUSE 3** | **MOUSE 4** | **MOUSE 5** | **MOUSE 6** | **MOUSE 7** | **MOUSE 8** |
| Number of mutants | 59 | 37 | 133 | 34 | 15 | 4 | 28 | 25 |
| Mutant frequency | $1.8 \times 10^{-6}$ | $7.3 \times 10^{-7}$ | $1.9 \times 10^{-6}$ | $6.2 \times 10^{-7}$ | $6.5 \times 10^{-7}$ | $2.0 \times 10^{-7}$ | $9.6 \times 10^{-7}$ | $4.1 \times 10^{-7}$ |

| | INOCULUM | LUNGS | KIDNEYS | BRAIN | ALL ORGANS |
|---|---|---|---|---|---|
| **MEDIAN FREQUENCY** | $5.6 \times 10^{-7}$ | $2.5 \times 10^{-6}$ | $5.5 \times 10^{-7}$ | $<7.4 \times 10^{-7}$ | $6.9 \times 10^{-7}$ |

**Table S3.** TE insertions in *URA3*/*URA5* of 5FOA-resistant mutants recovered from inoculum cultures and mouse organs.

| INOCULUM CULTURE | Culture 1 | Culture 2 | Culture 3 | Culture 4 | Culture 5 | Culture 6 | Culture 7 | | TOTAL |
|---|---|---|---|---|---|---|---|---|---|
| Mutants with TE insertions | 23 | 10 | 8 | 0 | 0 | 3 | 12 | | 56 |
| Total mutants screened | 24 | 24 | 24 | 24 | 24 | 24 | 24 | | 168 |
| Unique TE insertions | 2 | 2 | 1 | 0 | 0 | 1 | 1 | | 7 |

| LUNGS | Mouse 1 | Mouse 2 | Mouse 3 | Mouse 4 | Mouse 5 | Mouse 6 | Mouse 7 | Mouse 8 | TOTAL |
|---|---|---|---|---|---|---|---|---|---|
| Mutants with TE insertions | 37 | 21 | 75 | 18 | 1 | 0 | 15 | 22 | 189 |
| Total mutants recovered | 37 | 22 | 78 | 28 | 1 | 0 | 15 | 22 | 203 |
| Unique TE insertions | 6 | 3 | 9 | 3 | 1 | 0 | 7 | 3 | 32 |
| **KIDNEYS** | | | | | | | | | |
| Mutants with TE insertions | 8 | 15 | 53 | 1 | 14 | 2 | 4 | 0 | 97 |
| Total mutants recovered | 22 | 15 | 55 | 6 | 14 | 4 | 12 | 0 | 128 |
| Number of unique TE insertions | 3 | 2 | 4 | 1 | 2 | 2 | 2 | 0 | 16 |
| **BRAIN** | | | | | | | | | |
| Mutants with TE insertions | 0 | 0 | 0 | 0 | 0 | 0 | 0 | 3 | 3 |
| Total mutants recovered | 0 | 0 | 0 | 0 | 0 | 0 | 1 | 3 | 4 |
| Number of unique TE insertions | 0 | 0 | 0 | 0 | 0 | 0 | 0 | 1 | 1 |

| ALL ORGANS | Mouse 1 | Mouse 2 | Mouse 3 | Mouse 4 | Mouse 5 | Mouse 6 | Mouse 7 | Mouse 8 | TOTAL |
|---|---|---|---|---|---|---|---|---|---|
| Mutants with TE insertions | 45 | 36 | 128 | 19 | 15 | 2 | 19 | 25 | 289 |
| Total mutants recovered | 59 | 37 | 133 | 34 | 15 | 4 | 28 | 25 | 335 |
| Number of unique TE insertions* | 9 | 5 | 9 | 4 | 3 | 2 | 8 | 3 | 43 |

*Identical mutations in organs from the same mouse were counted as a single unique mutation. TCN12 (full length and solo LTR) insertions with the same insertion site and orientation were counted as a single unique mutation.

**Table S4.** Drug-resistance rates and proportions of TE insertions into target genes.

| Strain | Growth | Drug-resistance rate x $10^{-8}$ (95% CI) | Gene | T1 | TCN12 | T2 | T3 | CIRT1 | Rnd-3 | Unknown | Total TE | Total mutants screened |
|---|---|---|---|---|---|---|---|---|---|---|---|---|
| XL280 | 30º | **5FOA** 1.89 (0.94-2.75) | *URA3* | 2 | 0 | 0 | 0 | 0 | 0 | 0 | 2 | 90 |
| | | | *URA5* | 2 | 0 | 0 | 0 | 0 | 0 | 0 | 2 | |
| | 37º | 48 (39-62) | *URA3* | 60 | 1 | 0 | 0 | 0 | 0 | 0 | 61 | 251 |
| | | | *URA5* | 29 | 96 | 0 | 0 | 0 | 0 | 0 | 125 | |
| | 30º | **Rapamycin/FK506** 5.56 (4.39-12) | *FRR1* | 14 | 0 | 1 | 0 | 0 | 0 | 0 | 14 | 43 |
| | 37º | 43.5 (32.2-57.4) | *FRR1* | 35 | 15 | 1 | 0 | 0 | 0 | 0 | 51 | 61 |
| XL280 *rdp1*Δ | 30º | **5FOA** 4.62 (2.97-7.42) | *URA3* | 5 | 0 | 0 | 0 | 0 | 0 | 0 | 5 | 54 |
| | | | *URA5* | 18 | 0 | 0 | 1 | 0 | 0 | 2 | 21 | |
| | 37º | 54.9 (42.3-74.9) | *URA3* | 15 | 0 | 0 | 4 | 0 | 0 | 0 | 19 | 67 |
| | | | *URA5* | 11 | 26 | 0 | 1 | 0 | 0 | 0 | 37 | |
| NIH12 | 30º | 3.6 (2.76-5.07) | *URA3* | 0 | 0 | 0 | 2 | 1 | 0 | 0 | 3 | 62 |
| | | | *URA5* | 1 | 2 | 1 | 4 | 1 | 0 | 0 | 9 | |
| | 37º | 21.2 (16.5-23.6) | *URA3* | 0 | 1 | 0 | 8 | 1 | 0 | 0 | 10 | 91 |
| | | | *URA5* | 3 | 18 | 0 | 12 | 0 | 0 | 0 | 33 | |
| NIH433 | 30º | 2.07 (1.29-3.68) | *URA3/5* | 0 | 0 | 0 | 0 | 0 | 0 | 0 | 0 | 65 |
| | 37º | 12.9 (8.41-18.6) | *URA3/5* | 0 | 0 | 0 | 0 | 0 | 0 | 0 | 0 | 95 |
| 528 | 30º | 4.63 (2.0-5.4) | *URA3* | 1 | 0 | 0 | 8 | 0 | 0 | 0 | 9 | 57 |
| | | | *URA5* | 3 | 0 | 0 | 0 | 0 | 0 | 0 | 3 | |
| | 37º | 18.1 (11-23.4) | *URA3* | 8 | 0 | 0 | 12 | 0 | 4 | 0 | 24 | 76 |
| | | | *URA5* | 1 | 0 | 0 | 2 | 0 | 2 | 0 | 5 | |
| AD7-017 | 30º | 3.93 (1.51-6.59) | *URA3* | 0 | 0 | 0 | 0 | 0 | 0 | 0 | 0 | 44 |
| | | | *URA5* | 0 | 0 | 0 | 0 | 0 | 0 | 3 | 3 | |
| | 37º | 4.53 (3.49-5.65) | *URA3* | 0 | 0 | 0 | 0 | 0 | 0 | 0 | 0 | 42 |
